# MIRit: an integrative R framework for the identification of impaired miRNA-mRNA regulatory networks in complex diseases

**DOI:** 10.1101/2023.11.24.568528

**Authors:** Jacopo Ronchi, Maria Foti

## Abstract

**Background:** MicroRNAs are crucial regulators of gene expression that participate in nearly every cellular process. Due to their central role, miRNAs are frequently implicated in the development of pathological conditions, as their dysregulation significantly disrupts normal cellular functioning. Consequently, a thorough characterization of the involvement of dysregulated miRNAs in disturbed pathways is imperative for understanding the mechanisms behind diseases. Nevertheless, the limited availability of analytical methods for joint multi-omic analysis of miRNA-mRNA datasets frequently yields inconclusive and contradictory findings. Specifically, the lack of statistical frameworks designed for integrated analysis of miRNA and mRNA data across biological conditions, as well as the insufficiency of methods suitable for non-sample-matched data, severely restricts the understanding of miRNA networks and the reproducibility of the conclusions drawn. To address these limitations, here we present MIRit, an open-source and all-in-one R package that enables integrative multi-omic miRNA analyses using various statistical approaches.

**Results:** To showcase MIRit’s capabilities, we evaluated the pipeline on a thyroid cancer dataset, comparing 8 papillary thyroid carcinoma samples to contralateral healthy tissue, as well as on Alzheimer’s disease data, comparing temporal cortex samples from patients to those from healthy individuals. In the first case, MIRit revealed that upregulation of miR-146b-5p and miR-146b-3p caused downregulation of PAX8, resulting in decreased thyroid hormone transcription. In contrast, in the second dataset, MIRit showed that disrupted miRNA-mRNA networks in patients with Alzheimer’s disease impact neuroinflammation, glutamatergic signaling, and neuroprotection. In particular, overexpression of miR-320a-3p may reduce SERPINF1, potentially leading to the accumulation of *Aβ*_1*−*42_ fragments.

**Conclusions:** The adoption of MIRit’s pipeline permits a comprehensive evaluation of perturbed regulatory networks in human diseases through a novel approach for integrative pathway analysis of miRNA-mRNA data. Specifically, MIRit enables the characterization of impaired networks at the molecular level, providing an outstanding advantage in the functional characterization of key dysregulated factors involved in disease pathogenesis and progression. In conclusion, our findings demonstrate the efficacy and usability of MIRit, making it a valuable tool for researchers in the field.

## Background

MicroRNAs (miRNAs) are a class of small non-coding RNAs with an average length of 22 nucleotides, whose role is mainly to negatively regulate gene expression at a post-transcriptional stage. After their biogenesis, they typically promote mRNA degradation and translation inhibition by interacting with the 3’ untranslated region (3’ UTR) of target genes (1). Over the last few decades, evidence has progressively shown the involvement of miRNAs in mammalian pathways. Currently, it is estimated that more than 60% of human genes are controlled by miRNAs (2). Given their extensive regulatory activity, unbalanced miRNA expression can have dramatic consequences on cellular activities, and their dysregulation has been associated with several diseases (3). For example, the downregulation of miR-122, a liver-specific miRNA, has been shown to increase the proliferation and invasion of liver cells, which subsequently leads to the development of hepatocellular carcinoma (HCC) (4). Another example of the contribution of miRNAs to disease pathogenesis is miR-21, an oncogenic miRNA that plays a pivotal role in the onset and progression of several cancers, including glioblastoma and gastric cancer (5, 6). In light of these considerations, there has been a growing interest in the study of miRNAs with the aim of characterizing the compromised miRNA regulatory networks involved in human diseases. In this context, several approaches are available to estimate miRNA abundance at the omic level, including microarrays and RNA-Seq. Consequently, the determination of differentially expressed miRNAs (DE-miRNAs) in different biological conditions can be successfully performed using the same analytical procedures employed for transcriptomics. However, elucidating the biological consequences of dysregulated miRNAs is extremely difficult even after their identification. In fact, the limited functional annotation of miRNAs hinders our comprehension of miRNA-driven mechanisms in complex diseases, likely due to the ambiguity of miRNA-target gene interactions across various cell types and conditions. Nonetheless, using miRNA-mRNA multi-omic analyses offers a promising approach to exploring the biological effects of miRNA dysregulation and estimating the functional impact of impaired miRNAs. Indeed, using both miRNA and gene expression levels, we may be able to quantify the impact of each miRNA on the expression of its targets and reconstruct the disrupted molecular networks that may trigger the onset or exacerbation of a pathological state.

Despite the advantages of integrative miRNA-mRNA analyses, current approaches are rarely successful and poorly reproducible, limiting the ability to investigate the role of miRNA networks in disease pathogenesis. In particular, miRNA-mRNA analyses are dramatically affected by the choice of miRNA target genes. In this respect, the use of different prediction algorithms can have a drastic effect on the results of a study and thus limit the reproducibility of the conclusions drawn. Furthermore, the statistical methods used to associate miRNA and gene expression often lack rigor and accuracy. In this context, the most widely used approach is correlation analysis, which allows the statistical evaluation of the influence of miRNAs on gene expression. However, the statistical power to detect meaningful interactions is often reduced by incorrect implementations. Moreover, the use of correlation approaches drastically limits the availability of datasets for integrative analyses, as these methods require sample-matched miRNA and mRNA expression measurements, which are usually quite rare. Therefore, the lack of statistical approaches suitable for unpaired data represents a major bottleneck for integrative miRNA analyses. In addition, after identifying the dysregulated miRNA-target interactions that are involved in specific conditions, researchers may wish to explore the topological impact of such pairs on biological pathways and thereby identify the cellular functions that are most affected in the conditions of interest. Unfortunately, this type of integrative pathway analysis is often neglected, which hinders the interpretability of miRNA-mRNA analyses.

While state-of-the-art tools for integrative miRNA analysis, such as MiRComb (7), miRLAB (8), and anamiR (9), offer effective measures to estimate the impact of disease-associated miRNAs on the expression levels of their target genes, almost all approaches suffer from several common drawbacks. First, most methods use a combination of multiple databases to define miRNA targets, which often reports an unreasonably high number of interactions for each miRNA, with an inflated rate of false-positive hits. As a result, the vast number of retrieved interactions complicates downstream analysis, increases the burden of multiple testing, and limits the interpretation and experimental validation of reported biological results. Second, almost all tools rely solely on correlation analysis, which is not suitable for datasets without sample-matched measurements. In addition, some of the statistical approaches offered are not always appropriate for miRNA and mRNA datasets. For example, among the many statistical methods used in miRLAB to quantify the influence of miRNAs on target gene expression, the use of lasso and elastic-net regularized models is not recommended when the number of miRNAs targeting a gene is higher than the number of samples, which can be the case for some central hub genes. Finally, to the best of our knowledge, there is no tool available to perform an integrative pathway analysis that considers the topology of miRNA-gene interactions within biological pathways.

To address all these issues and fill these gaps, we developed MIRit, an open-source, comprehensive and all-in-one R framework that encompasses all the steps required to perform an efficient and statistically rigorous integrative miRNA-mRNA analysis. Moreover, compared to other tools, MIRit has multi-species support (Supplementary Table 1) and is designed to work with data derived from microarrays, RNA-Seq, miRNA-Seq, proteomics, and single-cell transcriptomics. In addition, MIRit implements novel statistical approaches that allow the analysis of datasets even without matched samples. In this paper, we describe the implementation of MIRit and demonstrate how it can be used in the context of human diseases. MIRit is freely available at https://github.com/jacopo-ronchi/MIRit.

## Implementation

### MIRit’s pipeline

To comprehensively examine the role of miRNAs in disturbed biological pathways, MIRit offers a versatile and robust workflow for executing integrated miRNA-mRNA multi-omic analyses from beginning to end. Specifically, MIRit’s sequential pipeline comprises multiple modules: 1) differential expression analysis, which identifies miRNAs and genes that vary across biological conditions; 2) functional enrichment analysis, which allows understanding the consequences of gene dysregulation; 3) single nucleotide polymorphism (SNP) association, which links miRNA dysregulations to disease-associated SNPs that occur at miRNA gene loci; 4) Target identification, which retrieves predicted and validated miRNA-target interactions through innovative state-of-the-art approaches that limit false discoveries; 5) Expression level integration, which relates the expression levels of miRNAs and mRNAs for both paired and unpaired data; 6) Topological analysis, which implements a novel approach to identify the perturbed molecular networks affecting biological pathways retrieved from KEGG (10), Reactome (11), and WikiPathways (12). An illustration of the general MIRit pipeline can be found in Figure 1.

**Fig. 1.**
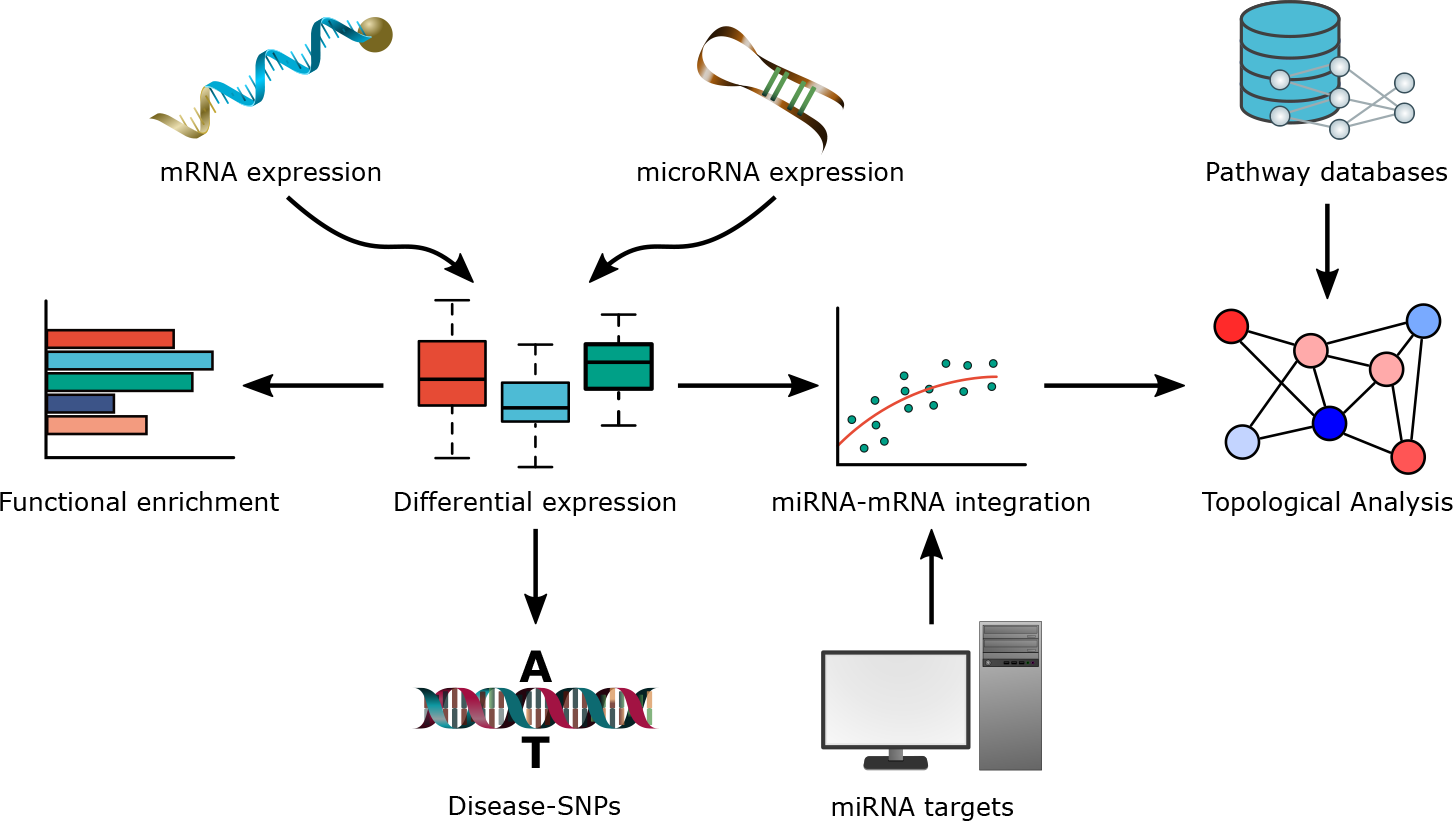
The MIRit pipeline. The workflow used by MIRit requires both miRNA and mRNA expression levels as input. Later, differential expression analysis is carried out for both miRNAs and genes to identify differentially expressed features, followed by gene functional enrichment to infer compromised biological processes. Additionally, we assess the presence of disease-associated SNPs within miRNA loci. Furthermore, we retrieve miRNA target genes from online databases and integrate their expression levels with those of miRNAs using different procedures for paired and unpaired datasets. Finally, MIRit performs a topological analysis to identify the impaired miRNA-mRNA regulatory networks that are responsible for the perturbation of biological pathways.

### Data preparation

As input, MIRit necessitates matrices of miRNA and gene expression measurements featuring samples as columns and genes/miRNAs as rows. The miRNA expression matrix’s row names should use miR-Base (13) nomenclature (e.g. hsa-miR-151a-5p, hsa-miR-21-5p), whereas the gene expression matrix’s row names must contain gene symbols conforming to HGNC (e.g. TYK2, BDNF, NTRK2). These matrices can accommodate multiple technologies and handle different types of values. For microarray studies, the expression matrices must be logarithmically transformed and normalized. This can be accomplished by using the Robust Multi-array Average (RMA) algorithm within the oligo R package (14) or through other quantile normalization strategies. On the other hand, for RNA-Seq and miRNA-Seq experiments, a simple count matrix must be provided. At this stage, MIRit saves expression matrices and sample metadata in an object of class MirnaExperiment, which extends the MultiAssayExperiment class from the homonym package (15) to accommodate genomic data of both miRNAs and their targets. In particular, this object is capable of storing information on expression levels, differential expression outcomes, miRNA-target pairs, and miRNA-gene integration analysis. MIRit’s pipeline employs this object for all further analysis.

### Differential expression analysis

After setting up expression matrices, differential expression analysis is performed for both miRNAs and genes. In this regard, MIRit offers several options based on the technology used to produce expression data. When expression values are obtained through microarrays, MIRit uses the standard pipeline in the limma package (16) to estimate differentially expressed features. On the other hand, when deriving expression values from RNA-Seq or miRNA-Seq experiments, MIRit offers a choice between various approaches, such as edgeR (17), DESeq2 (18), and limma-voom (19). Importantly, MIRit offers the ability to fully customize the parameters used for differential expression analysis, which allows for more fine-grained control and facilitates the adoption of strategies different from the standard pipelines proposed in these packages. In this concern, MIRit is appropriate for various analysis types, as it supports complex and multivariate experimental designs. Additionally, users can independently conduct and load their own analyses. This enables MIRit to process results from a range of tools and technologies, including proteomic data analyzed using specialized R packages such as MSqRob2 (20). Lastly, MIRit offers numerous functions for displaying differential expression results. These encompass volcano plots, bar plots, box plots, and violin plots.

### Functional enrichment of dysregulated genes

After performing differential expression analysis, it is common to obtain extensive lists of features whose expression changes between biological conditions. Nonetheless, such lists typically lack knowledge and comprehension as to which biological functions are perturbed in the conducted experiments. Therefore, various approaches are typically used to determine which cellular processes are dysregulated under the given conditions of interest. Among these methods, the initial approach developed is referred to as over-representation analysis or ORA (21), for short. Its objective is to establish if genes identified and annotated to individual gene sets, such as biological pathways or ontological terms, exhibit differential expression exceeding what would be anticipated randomly. To accomplish this, p-values are computed for each gene set through the hypergeometric distribution. Then, following multiple testing adjustments, significant categories represent the biological processes that vary across the tested conditions. In addition, an alternative approach called gene set enrichment analysis (GSEA) (22) is also available. The GSEA algorithm takes as input a list of genes ranked according to a particular criterion, and then iterates down the list to assess whether members of a given gene set are normally distributed, or clustered at the top or bottom of the list. Unlike ORA, which is limited in detecting small differences in gene expression, GSEA is capable of identifying categories with genes that exhibit coordinated but subtle changes. Although GSEA is a widely-used approach for functional enrichment, Wu and Smyth (23) demonstrated that inter-gene correlations can affect the reliability of functional enrichment analyses. To resolve this issue, they developed the Correlation Adjusted MEan RAnk gene set test (CAMERA), another competitive method used for gene functional enrichment analysis. The primary benefit of this approach is its ability to adapt the gene set test statistic based on inter-gene correlations.

MIRit provides extensive support for functional enrichment of genes using ORA, GSEA, and CAMERA, empowering the user to deduce compromised biological functions using the preferred method. Notably, for ORA, MIRit performs the enrichment separately for upregulated and downregulated genes, an approach that has been shown to be more powerful than enriching all differentially expressed genes (DEGs) (24). Additionally, MIRit uses the geneset R package (25), which provides access to numerous biological databases. This feature allows MIRit to perform functional enrichment analyses on gene sets from sources such as Gene Ontology (GO) (26), KEGG, WikiPathway, MsigDb (22), Reactome, MeSH, DisGeNET (27), Disease Ontology (28), Network of Cancer Genes (29), and COVID-19 (30). Moreover, MIRit features intuitive functions such as bar plots, dot plots, and GSEA plots to simplify the visualization of enriched categories.

### Association of miRNAs with disease-SNPs

Further-more, MIRit provides a useful tool for examining the overlap between DE-miRNA loci and genomic variants linked to human illnesses. This is important because SNPs present within miRNA gene loci can have dramatic impacts on the biological functioning of these transcripts. In fact, a SNP found within a miRNA gene could modify its expression or alter the spectrum of miRNA targets, leading to potential pathogenetic mechanisms. In detail, MIRit retrieves disease-associated SNPs using the gwasrapidd package (31) which directly queries the NHGRI-EBI Catalog of published genome-wide association studies (32). Next, the tool obtains the genomic positions of DE-miRNAs from the Ensembl database (33) and overlays them with the positions of disease-associated SNPs. Finally, disease-associated SNPs found within miRNA genes are reported.

### MiRNA targets retrieval

Before conducting integrative miRNA-mRNA analyses, it is essential to determine the target genes of DE-miRNAs to examine if their expression levels are influenced. Several resources have been developed to predict and collect miRNA-target interactions over the years. These can be categorized into two types: prediction databases, which contain computationally determined information about miRNA-target interactions, and validated databases, which only contain interactions that have been proven through biomolecular experiments. The choice of the resource to use for miRNA target identification will significantly affect the analysis outcome. In this context, several bioinformatic pipelines prioritize validated interactions, even though they tend to be less numerous than predicted pairs. Conversely, predicted pairs are more abundant but have a higher incidence of false positives. MiRNA target prediction algorithms are also hindered by a low degree of overlap across different tools. To tackle this challenge, a number of ensemble methods have been devised to combine predictions made by different algorithms. Initially, miRNA-target pairs were considered significant if predicted by multiple tools (intersection method). However, this approach may not capture a noteworthy number of significant associations. On the other hand, other approaches merged predictions from a variety of algorithms (union method). Although the union method identifies more true relationships, it also results in a higher proportion of false discoveries. As a consequence, other ensemble methods have started to incorporate alternative statistics to rank miRNA-target predictions obtained from multiple algorithms. Among the recently developed ensemble methods, the microRNA Data Integration Portal (mirDIP) database (34) stands out as a top performer. It combines miRNA target predictions from 24 different resources by using an integrated score derived from different prediction metrics. In this way, mirDIP reports more accurate predictions compared to those of individual tools (34, 35).

MIRit predicts miRNA-target interactions using the mirDIP database (version 5.2) and further improves accuracy by merging miRNA-target pairs obtained from mirDIP with experimentally validated interactions in the miRTarBase database (version 9) (36).

### Investigation of the impact of miRNA expression changes on target genes

After defining the targets of DE-miRNAs, an integrative analysis of miRNA and gene expression levels can be performed. This analysis is useful as it enables the consideration of putative miRNA-target pairs only when an inverse relationship is observed in the data. As previously mentioned, MIRit can work with both paired and unpaired data by using different statistical approaches, such as correlation analysis, association tests, and rotation gene-set tests.

When both miRNA and gene expression measurements are available for the same samples, a correlation analysis is the recommended procedure. In statistics, correlation is a measure that expresses the extent to which two random variables are dependent. In this case, the aim is to evaluate if a statistical relationship exists between the expression of a miRNA and its targets’ expression. To determine the degree of correlation, several statistical coefficients can be used. Among them, the most frequently employed are Pearson’s *r*, Spearman’s *ρ*, and Kendall’s *τ*_*b*_. Pearson’s *r* is the most widely used for determining the association between miRNA and gene expression; however, it assumes a linear relationship. This is typically not true for miRNAs, as their interactions with targets are characterized by imperfect complementarity. In addition, miRNAs can target multiple genes with different binding sites, suggesting that a simple linear relationship may not adequately describe the complexity of these interactions. In contrast, Spearman’s and Kendall’s Tau-b correlation coefficients are more appropriate for representing the interplay between miRNAs and target genes because they are robust to non-linear relationships and outliers. Nevertheless, Kendall’s correlation relies solely on the number of concordant and discordant pairs and is less sensitive than Spearman’s correlation. Therefore, when numerous ties are present or the sample size is small, Kendall’s correlation may have lower detection power. This is why Spearman’s correlation coefficient is the default measure used in MIRit to evaluate the correlation between miRNA and gene expression. Moreover, as miRNAs primarily function as negative regulators, only negatively correlated miRNA-target pairs are considered. Then, statistical significance is estimated by a one-tailed t-test. Furthermore, to address potential batch effects in expression matrices prior to correlation analysis, MIRit provides an option to remove the variability due to unwanted technical artifacts. This involves fitting a linear model to the expression data, then adjusting the matrices by subtracting the variation due to batch effects. This is necessary because correlation analysis evaluates the concordance of measurements taken on a per-sample basis, which can be confounded by technical variations. Importantly, the resulting batch effect-free matrices cannot be used for differential expression analysis. For this purpose, it is recommended to model the batch variables in the linear model to avoid underestimating the residual degrees of freedom, which could lead to overoptimistic standard errors, t-statistics, and p-values.

However, it is impossible to directly measure the impact of miRNA expression on the levels of its targets for unpaired data because miRNA and gene measurements lack sample correspondence. In such cases, one-sided association tests can be used to determine whether the targets of down-regulated miRNAs are statistically enriched in up-regulated genes, and vice versa, to assess if the targets of up-regulated miRNAs are statistically enriched in down-regulated genes. In order to evaluate the impact of DE-miRNAs on their target genes, MIRit has the capability to use two distinct one-sided association tests: Fisher’s exact test and Boschloo’s exact test. These tests involve a statistical procedure that estimates the association between two dichotomous categorical variables. In this case, the aim is to evaluate if there is a notable discrepancy in target proportion between differentially and non-differentially expressed genes for each miRNA. To achieve this, a 2x2 contingency table is constructed for each miRNA as demonstrated in Table 1.

**Table 1.**
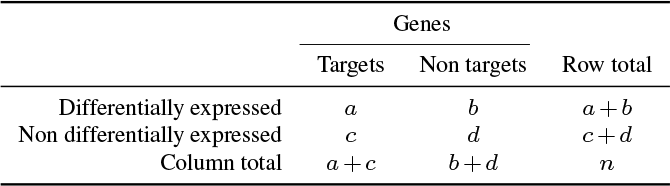
The 2x2 contingency table that MIRit uses for one-sided association tests.

Once the contingency table has been defined, the p-value for the Fisher’s exact test can be calculated using the Equation 1:

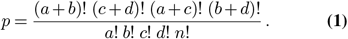

MIRit manages the entire process and calculates p-values for each DE-miRNA. Fisher’s p-values are subsequently adjusted for multiple testing and significant findings are reported. Moreover, it is possible to estimate Fisher’s p-values using Lancaster’s mid-p correction, as it has been demonstrated to increase statistical power while preserving type-I error rates (37).

However, Fisher’s exact test is a conditional test that requires both the rows and columns sums of a contingency table to be fixed. Nevertheless, for genomic data, it is likely that different datasets could result in a varying number of DEGs. Hence, MIRit defaults to using a variant of Barnard’s exact test, named Boschloo’s exact test, which is more suitable for contingency tables with variable group sizes. Furthermore, it is possible to demonstrate that Boschloo’s test is uniformly more powerful than Fisher’s exact test (38).

Lastly, unpaired data can take advantage of linear models and rotation gene-set tests to estimate the impact of DE-miRNAs on target gene expression. The purpose of this approach is to assess whether each miRNA leads to differentially expressed targets in the opposite direction. In particular, ‘fry’, a fast approximation of rotation gene-set testing (39) implemented in the limma package, can be employed to statistically measure the impact of miRNAs on changes in expression of their target genes.

### Topology-Aware Integrative Pathway Analysis (TAIPA)

Once the dysregulated miRNA-target pairs have been identified, the typical goal is to infer the altered cellular processes and functions. Procedures that aspire to attribute significance to lists of dysregulated factors fall under the name pathway analysis (PA). Over the years, various types of PA have been developed. The first-generation methods, which include the aforementioned ORA, rely exclusively on the proportion of differentially expressed molecules. On the other hand, the second-generation methods, known as functional class scoring (FCS), include GSEA, among others. In this analysis, different scoring statistics are used to estimate the enrichment of each gene set. The primary advantage of second-generation methods is their sensitivity to small variations as they extract information from all studied molecules. However, both first and second-generation methods neglect the topology of pathway networks, disregarding the direction of biological interactions. To overcome this limitation, a third generation of methods, known as Topology-Based, has been introduced to take into account the topology of biological networks (40).

One interesting approach in this context is to incorporate significant miRNA-mRNA interactions into biological pathways obtained from pathway databases, to use topology-based PA methods to deduce dysregulated miRNA-mRNA regulatory networks. The commonly employed methods in this category include SPIA (41), PRS (42), CePa (43), TAPPA (44), TopologyGSA (45), Clipper (46), and DEGraph (47). However, TAPPA, TopologyGSA, Clipper, and DEGraph do not use fold changes as node attributes, making them unsuitable for datasets without sample-matched expression (48). Conversely, while PRS and CePa only require fold changes to calculate pathway scores, they do not take into account the direction and concordance of biological interactions, which are critical for understanding perturbations in miRNA-mRNA networks. On the other hand, SPIA considers interaction direction and has been proposed by Nguyen et al. for integrative miRNA-mRNA analyses to overcome the bottleneck of sample-matched datasets (49). Nonetheless, SPIA inherently overweights the contribution of root nodes, which in this case consist of dysregulated miRNAs that inversely correlate with target genes (48). As a result, the application of SPIA to miRNA-augmented pathways may lead to an over-estimation of pathway significance. Therefore, even though numerous topology-based approaches could be deemed fit for this objective, none of them prove to be suitable for integrative miRNA analyses. To address this issue, we developed a new approach named Topology-Aware Integrative Pathway Analysis (TAIPA), which specifically focuses on detecting altered molecular networks in miRNA-mRNA multiomic analyses by considering the topology of biological pathways and miRNA-target interactions.

Initially, MIRit employs the graphite R package (50) to obtain pathways from KEGG, Reactome, or WikiPathways. Next, significant miRNA-target pairs identified through correlation or association analyses are integrated into the biological pathways. Each miRNA-augmented network is denoted as a graph *G*(*V, E*), where *V* represents nodes and *E* represents the relationships between nodes. Then, nodes that are not significantly differentially expressed are assigned a weight *w*_*i*_ = 1, whereas differentially expressed nodes are assigned a weight *w*_*i*_ = |Δ*E*_*i*_|, where Δ*E*_*i*_ is the linear fold change of the node. Moreover, to consider the biological interaction between two nodes, namely *i* and *j*, we define an interaction parameter *β*_*i→j*_ = 1 for activation interactions and *β*_*i→j*_ = *−*1 for repression interactions. Subsequently, the concordance coefficient *γ*_*i→j*_ is defined as described by Equation 2:

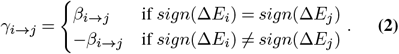

Later in the process, a breadth-first search (BFS) algorithm is applied to topologically sort pathway nodes so that each individual node occurs after all its upstream nodes. Nodes within cycles are considered leaf nodes. At this point, a node score *ϕ* is calculated for each pathway node *i* through Equation 3:

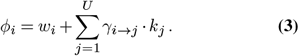

where *U* represents the number of upstream nodes, *γ*_*i→j*_ denotes the previously defined concordance coefficient, and *k*_*j*_ is a propagation factor defined in Equation 4, which represents the total network disturbance accumulated at node *j*.

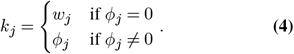

Finally, the pathway score Ψ is calculated according to Equation 5:

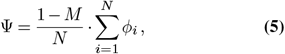

where *M* represents the proportion of miRNAs in the pathway, and *N* represents the total number of nodes in the network. In this context, it is essential to normalize the sum of node scores for both the proportion of miRNAs *M* and the overall node count *N*, because negatively correlated miRNA-target pairs always exhibit expression concordance.

Then, to compute the statistical significance of each pathway score, a permutation procedure is applied. In detail, for each pathway, the fold changes of miRNAs and genes are permuted a significant number of times and then the pathway score is calculated for each permutation. Moreover, miRNA and gene permutations are conducted separately because, unlike genes, miRNAs present in augmented pathways are uniquely differentially expressed. Hence, only DE-miRNAs are considered for miRNA permutations. Next, as with the PRS method, pathway scores undergo normalization following the method proposed by Tian et al. (51). In essence, to enable the comparison of pathway scores, each score is normalized on the basis of the specific null distribution of that pathway generated through random permutations. In particular, to obtain the standardized pathway score *Ψ*_*N*_, we subtract the mean score of the permuted sets 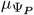 and then divide by the standard deviation of the permuted scores 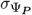, as per Equation 6:

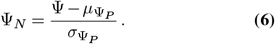

In the same way, permuted pathway scores Ψ_*P*_ are standardized according to Equation 7:

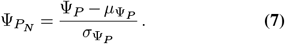

In the end, we calculate p-values for each pathway based on the proportion of permuted sets that reported a normalized pathway score 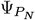 equal to or more extreme than the observed score *Ψ*_*N*_, as outlined in Equation 8:

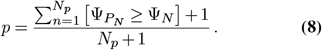

where *N*_*p*_ is the number of permutations, and the expression 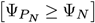 within Iverson brackets is equal to 1 if the statement is true and 0 otherwise. It is crucial to note that Equation 8 requires the addition of one to the numerator and the denominator to avoid p-values being equal to zero. According to Phipson and Smyth (52), the omission of this term results in a failure to control the type-I error rate, especially in scenarios with multiple comparisons, where p-values equal to zero would remain zero even when multiplied by a factor during multiple testing corrections.

Finally, p-values are either adjusted through the Benjamini-Hochberg procedure or through the max-T approach described by Westfall and Young (53), which could be more powerful in some scenarios. Regarding the entire method, increased permutations result in more stable p-value calculations, even though they require more time. Thus, a minimum of 10,000 permutations is recommended, with no fewer than 1,000. In this context, the computation of pathway scores for so many networks per pathway is computationally intensive. Therefore, pathway score calculation has been employed using the C++ language and parallel computing techniques have been adopted to decrease running times.

## Results and Discussion

To showcase the capabilities of MIRit in understanding human diseases, we examined two multi-omic datasets. First, we used MIRit to investigate how disrupted miRNA-mRNA networks affect thyroid functionality in a sample-matched papillary thyroid carcinoma dataset. Moreover, we present an example of how MIRit was applied to an unpaired dataset to explore the functional role of compromised miRNA regulatory networks in patients with Alzheimer’s disease.

### Example 1: papillary thyroid carcinoma (PTC)

In this example, we will analyze sequencing data obtained from Riesco-Eizaguirre et al. (54) who used RNA-Seq and miRNA-Seq to profile gene and miRNA expression in 8 papillary thyroid carcinoma tumors and corresponding contralateral tissue from the same patients. Initially, we downloaded raw count matrices from the Gene Expression Omnibus (GEO accession number: GSE63511) database (55) through the GEOquery R package (56). Importantly, only PTC samples with available matched gene expression measurements were included. To facilitate user accessibility, MIRit includes raw count matrices that allow for straightforward replication of this study’s results.

Then, we used multidimensional scaling (MDS) plots to examine variability in gene and miRNA expression. As a result, it emerges that disease condition causes well-separated samples for both genes (Figure 2a) and miRNAs (Figure 2b), indicating minimal impact of batch effects. Next, we adopted the edgeR pipeline implemented in MIRit to identify differentially expressed genes and miRNAs between PTC samples and healthy thyroid tissue. Specifically, since each individual was tested twice in this experimental design - once for cancer tissue and once for healthy contralateral thyroid - patient ID served as a covariate in the model to prevent individual differences from confounding disease-related alterations. Following this analysis, we identified 265 DEGs (Supplementary Table 2) and 40 DE-miRNAs (Supplementary Table 3), with a fold change greater than 2 and an FDR-adjusted p-value less than 0.05. As shown in Figures 2c and 2d, upregulated and downregulated genes are equally balanced, while miRNAs demonstrate consistently higher log2 fold changes. Importantly, Figures 2e-h show a significant downregulation of four genes critical to the normal functioning of the thyroid gland. Among these are thyroglobulin (TG), the thyroid hormone; thyroid peroxidase (TPO), an enzyme required for TG production; iodothyronine deiodinase 2 (DIO2), which activates TG; and Paired Box 8 (PAX8), a transcription factor necessary for TG transcription.

**Fig. 2.**
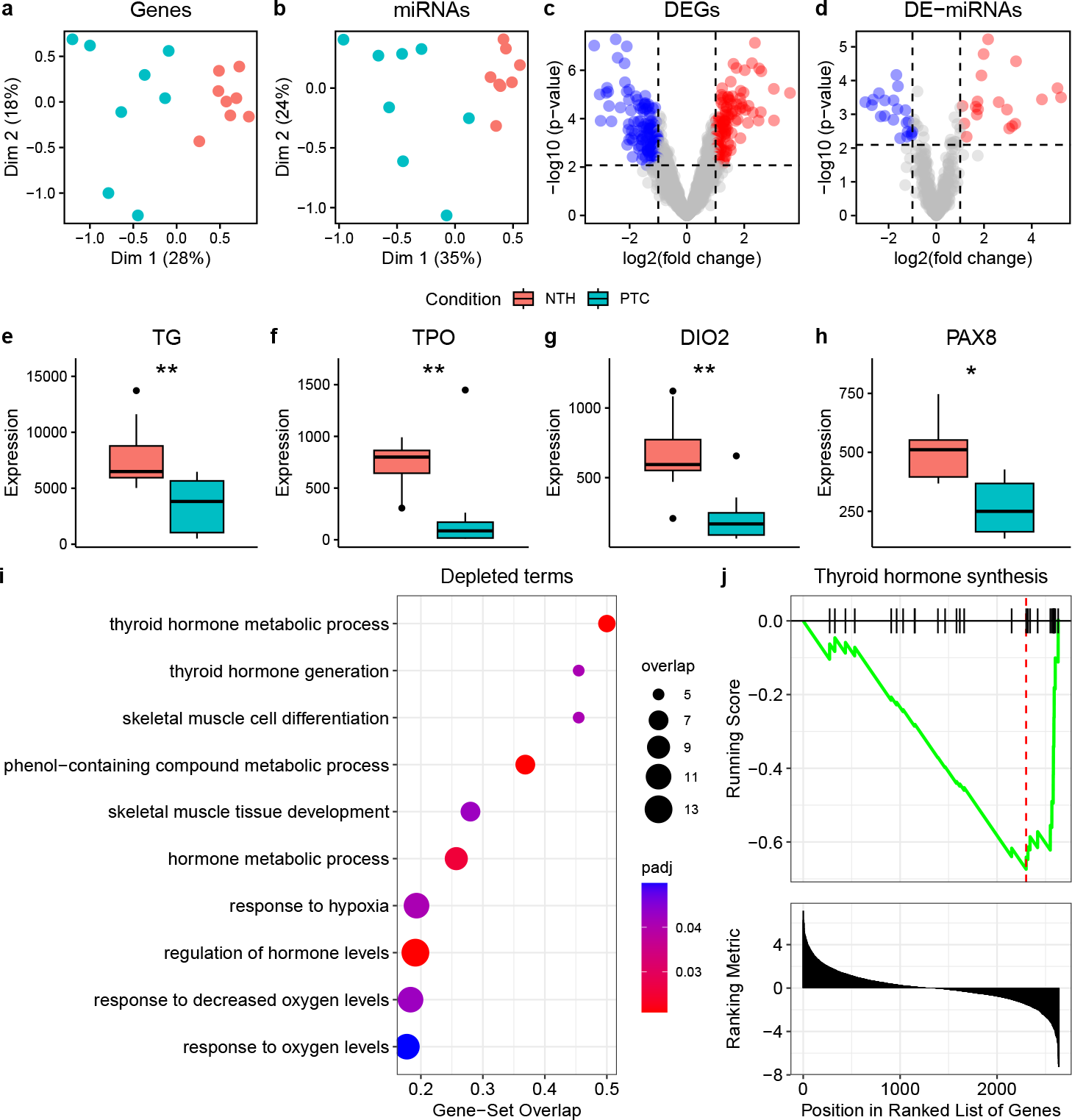
MiRNome and transcriptome analysis of PTC samples. **(a, b)** Expression similarities in the multidimensional space between PTC samples and normal thyroid (NTH), for both genes (a) and miRNAs (b), respectively. **(c, d)** Differences in gene (c) and miRNA (d) expression between PTC and NTH samples. **(e-h)** The decreased expression of several factors crucial to the normal functioning of the thyroid gland, namely thyroid hormone (TG), thyroid peroxidase (TPO), iodothyronine deiodinase 2 (DIO2), and Paired Box 8 (PAX8). **(i)** The enrichment of GO terms for downregulated genes, which confirms the impairment of thyroid hormone-related pathways. **(j)** The significant depletion identified by GSEA of the genes that belong to the Thyroid hormone synthesis pathway in the KEGG database.

Furthermore, MIRit was used to perform functional enrichment analysis of dysregulated genes. Initially, ORA was conducted on GO gene sets to investigate compromised functions in thyroid cancer, and categories with p-values below 0.05 after Benjamini-Hochberg correction were labeled as significant. As a result, it is possible to observe the depletion of genes accountable for thyroid hormone metabolism and generation, which highlights the impairment of this pathway in thyroid cancer (Figure 2i). Moreover, to assess the dysregulated networks, GSEA was conducted on human pathways obtained from the KEGG database. In agreement with ORA findings, GSEA indicated a significant decline in the “Thyroid hormone synthesis” pathway (FDR < 0.05), which is crucial for the adequate functioning of human thyroid (Figure 2j).

Subsequently, MIRit was employed to collect the targets of DE-miRNAs from mirDIP and miRTarBase. In detail, the top 5% of predicted interactions from mirDIP were considered along with validated pairs as true targets. As a result of this procedure, 13605 putative miRNA-target pairs were identified for the 40 DE-miRNAs. Subsequently, Spearman’s correlation analysis was performed to integrate miRNA and gene expression values. In this respect, 215 miRNA-target interactions showed a statistically significant inverse correlation, with a Spearman’s coefficient below -0.5 and an FDR-adjusted p-value lower than 0.05 (Supplementary Table 4). Of note, the correlation analysis demonstrated that miR-146b-5p, the most upregulated miRNA, displays an inverse correlation with DIO2, a critical factor in thyroid hormone function (Figure 3a). Furthermore, it has been demonstrated that miR-146b-5p and miR-146b-3p are negatively correlated with PAX8, which directly induces transcription of TG (Figures 3b and 3c). Additionally, functional analysis of the anti-correlated targets was performed using the Disease Ontology database with ORA enrichment (Figure 3d). As a result, when we enrich for downregulated targets, we observe a depletion of diseases associated with an overactive thyroid gland, such as goiter and hyperthyroidism (FDR < 0.05). This suggests the involvement of miRNAs in thyroid dysfunction.

**Fig. 3.**
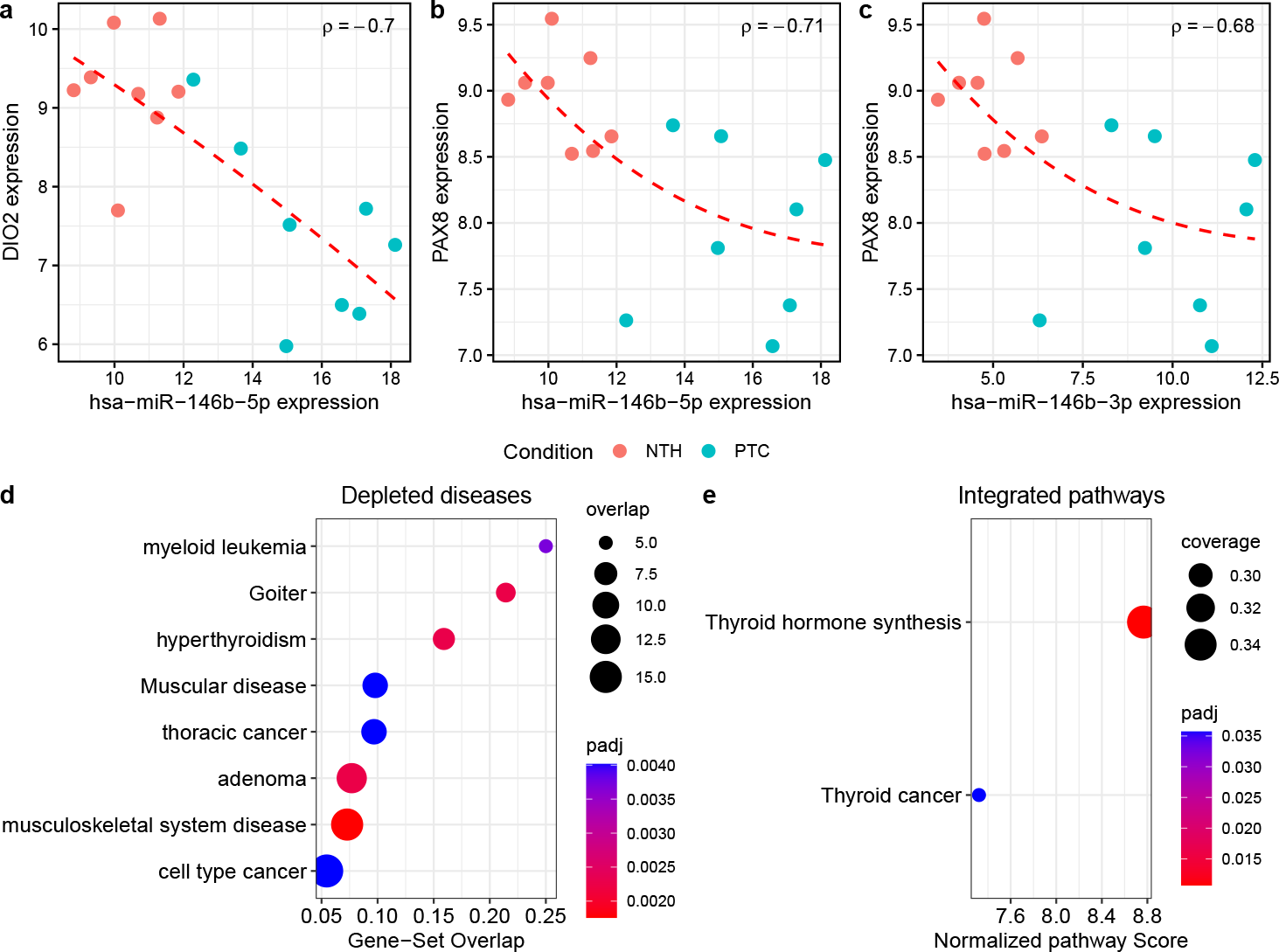
Integrative analysis of miRNA and gene expression levels. **(a-c)** The correlation existing between miR-146b-5p and DIO2, miR-146b-5p and PAX8, and miR-146b-3p and PAX8, respectively. From these charts it’s easy to note how the strong upregulation of these miRNAs in PTC samples is responsible for the concomitant downregulation of DIO2 and PAX8. **(d)** The ORA enrichment of downregulated genes that are negatively correlated with miRNA expression. This graph indicates how genes linked to conditions characterized by an overly active thyroid gland become depleted. **(e)** The significantly perturbed miRNA-amplified pathways as detected by TAIPA. The integrative miRNA-mRNA analysis is consistent with previous findings, hence emphasizing how miRNA and mRNA dysregulations in thyroid cancer ultimately result in severe thyroid malfunctioning.

Finally, we used MIRit to perform TAIPA and fully explore the impaired molecular networks in thyroid cancer. Firstly, we excluded 155 biological pathways with less than 10% of nodes with differential expression measurements. This step is necessary because, during differential expression analysis, lowly expressed features are removed. Subsequently, we performed integrative pathway analysis on biological networks from the KEGG database. As shown in Figure 3e, the pathways for thyroid hormone production and thyroid cancer processes are the most disrupted (max-T-adjusted p-value < 0.05). These results are consistent with our previous findings, which indicated thyroid gland dysfunction. At the molecular level, the increased expression of miR-146b-3p and miR-146b-5p is strongly associated with the downregulation of PAX8, which in turn leads to decreased thyroid hormone transcription, as shown in Figure 4. It is important to note that this phenomenon could have clinical significance, as it potentially contributes to the development of iodine resistance in thyroid cancer. This specific molecular mechanism revealed by MIRit has been confirmed through multiple in vitro assays (54), underscoring the utility of the presented pipeline for identifying perturbed molecular networks in human diseases.

**Fig. 4.**
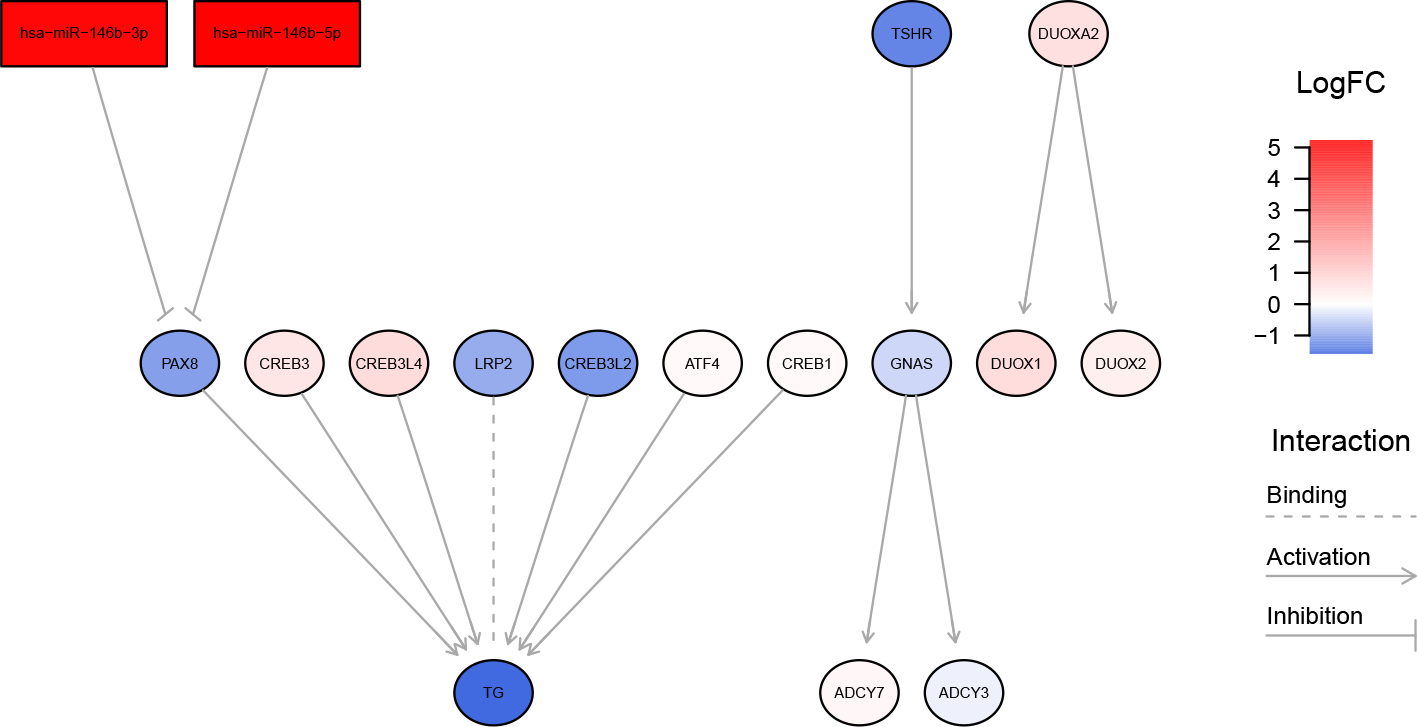
The impairment of thyroid hormone synthesis in thyroid cancer. TAIPA findings suggest that elevated levels of miR-146b-5p and miR-146b-3p in PTC samples are associated with decreased expression of PAX8, which leads to reduced thyroid hormone transcription.

### Example 2: Alzheimer’s disease

In our second case study, we demonstrate the abilities of MIRit in assessing the role of miRNA dysregulations in the prefrontal cortex of Alzheimer’s disease (AD) patients. To minimize expression variability and explore the network dysregulations specific to AD pathology, we restricted our analysis to studies focusing solely on Brodmann area 9 (BA9). With the intention of demonstrating MIRit’s utility in the evaluation of unpaired data, we selected two distinct studies. The initial dataset (GEO accession: GSE63501) comprises a small RNA-Seq experiment (57) that examines post-mortem BA9 tissues from 6 patients with AD, 3 subjects with tangle-predominant dementia (TPD) and 7 healthy controls. On the other hand, the second dataset (GEO accession: GSE150696) used microarray expression data of BA9 obtained through the Affymetrix Human Transcriptome Array 2.0, including samples from 9 patients with AD, 12 patients with dementia with Lewy bodies (DLB), 12 subjects with Parkinson’s disease dementia (PDD), and 9 healthy donors (58). For our analysis, we kept only the expression data of Alzheimer’s disease patients and healthy controls. Initially, the relevant FASTQ files for miRNA expression were downloaded from the Sequence Read Archive (SRA) (59) using the SRA Toolkit. Subsequently, we used the miRge 3.0 pipeline (60) to preprocess miRNA reads, which involves adapter trimming and quality control with Cutadapt (61), and alignment of reads against a modified miRNA library with bowtie (62). Ultimately, miRNA raw counts were determined following miR-Base nomenclature. For adapter trimming, we used TC-CGACGATC as the 5’ adapter sequence and TCGTATGC-CGTCTTCTGCTTGT for the 3’ adapter. Information on the number of sequenced reads and identified miRNAs for each sample can be found in Supplementary Table 5. Then, we acquired the transcriptomes of both AD and control samples by downloading microarray CEL files from GEO and importing them into R via the oligo package (14). Next, we used the RMA algorithm to normalize probe expression and summarize values at the transcript level. After annotating the probes, we removed features that had low expression levels, as depicted in Supplementary Figure 1. Later, we generated MDS plots for the miRNA and mRNA datasets to confirm AD mechanism-related transcriptional variation (Figures 5a and 5b). Following this, we used MIRit for differential expression analysis of both mRNAs and miRNAs. With regard to mRNAs, differential expression was assessed through the limma pipeline with estimated array quality weights. In this concern, age and sex were included in the linear model as covariates to account for additional biological variation. After this step, 353 DEGs (fold change > 1.5 and FDR-adjusted p-value < 0.05) were identified (Supplementary Table 6), as shown in Figure 5c. Moreover, we employed the limmavoom pipeline for identifying DE-miRNAs in AD patients. As a result, we have identified 12 DE-miRNAs (Supplementary Table 7) with an absolute fold change greater than 2 and a p-value less than 0.05 (Figure 5d). Subsequently, we used the `findMirnaSNPs` function in MIRit to identify potential AD-related SNPs occurring within the genomic loci of DE-miRNAs. Notably, we identified the presence of rs2632516, a variant linked with AD susceptibility, within the host gene of miR-142, one of the previously identified DE-miRNAs. Specifically, this polymorphism is located upstream of the miR-142 gene on the negative strand (Figure 5e), and may play a role in its transcriptional regulation. In this regard, Latini et al. showed that the presence of the rs2632516 variant allele in patients with systemic lupus erythematosus (SLE) is significantly associated with diminished miR-142 expression (63), which we found to be the second-most downregulated miRNA in the BA9 region of AD patients. However, additional research is necessary to uncover the functional significance of this polymorphism in the pathophysiology of Alzheimer’s disease.

**Fig. 5.**
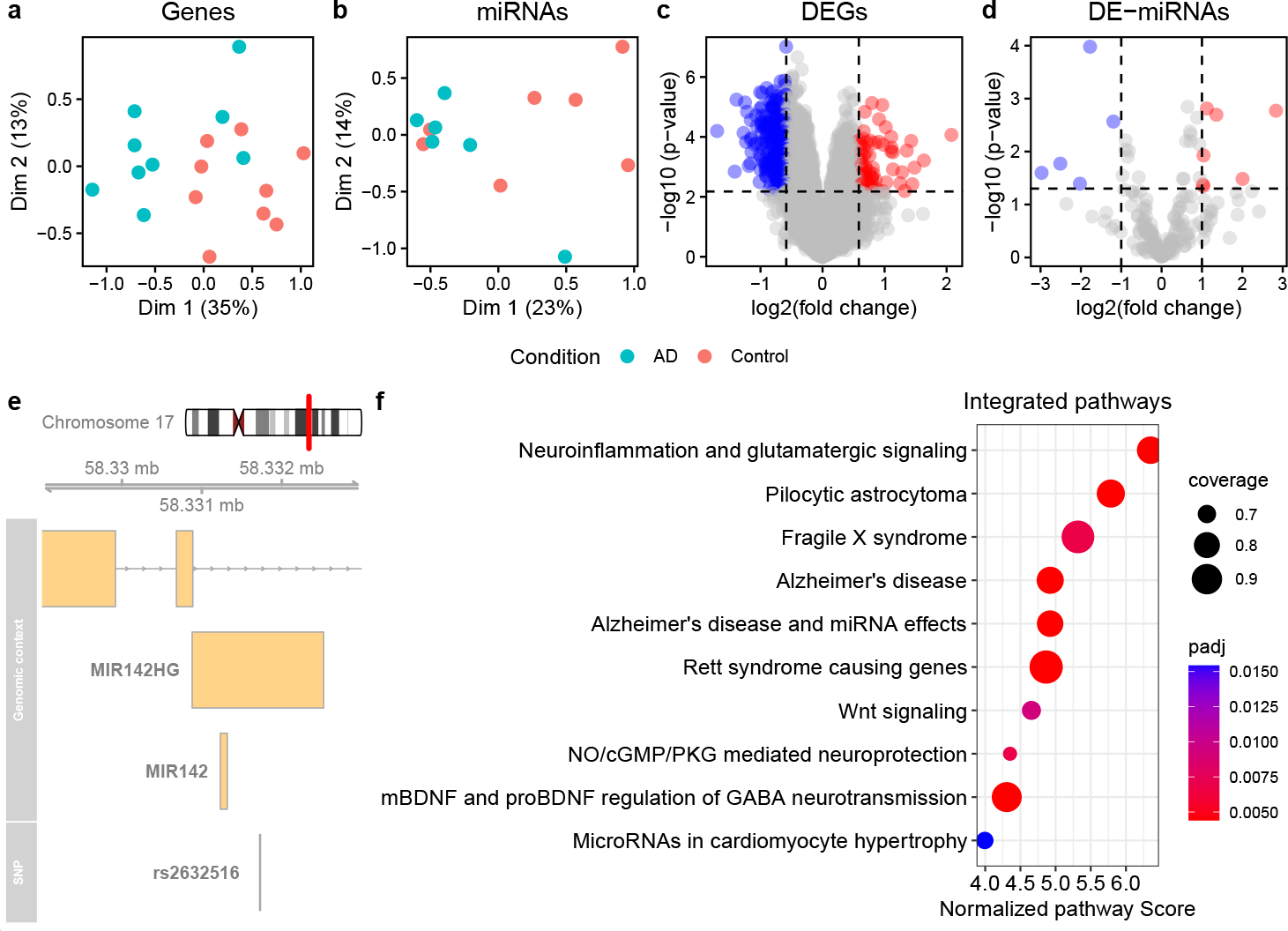
The role of altered miRNA expression in the BA9 region of patients with Alzheimer’s disease. **(a, b)** Transcriptional variability for genes (a) and miRNAs (b), respectively. **(c, d)** Results of differential expression analysis concerning genes (c) and miRNAs (d). **(e)** The presence of the rs2632516 variant in the miR-142 host gene on the negative strand, which may lead to impaired transcription of this miRNA. **(f)** TAIPA results reporting the disruption of multiple miRNA-mRNA networks relevant to AD pathophysiology, including neuroinflammation, glutamatergic signaling, synaptic plasticity, and neuroprotection.

Then, we collected miRNA targets using the top one-third of predicted interactions from the mirDIP database, along with validated pairs obtained from miRTarBase. Following this step, we retrieved 62748 putative miRNA-target pairs. Next, as this analysis relies on unpaired data, we performed Boschloo’s exact test to estimate the association between miRNA and mRNA expression changes. In the end, a statistically significant association was discovered between 7 miRNAs and 250 target genes, with an FDR-adjusted p-value below 0.05 (Supplementary Table 8). This number highlights the considerable impact of miRNA dysregulations on gene expression alterations in the prefrontal cortex of AD patients, revealing that over 70% of DEGs are targeted by at least one DE-miRNA. Finally, our aim was to investigate miRNA-mRNA network perturbations in BA9 region of individuals with AD. To achieve this, we performed TAIPA using biological networks obtained from the WikiPathways database. After excluding 68 pathways with less than 10% of nodes with available expression measurements, MIRit demonstrated the considerable impairment of various pathways that play a critical role in AD pathology (FDR-adjusted p-value < 0.05), as illustrated in Figure 5f. In addition to the pathways related to AD and miRNA effects in AD, TAIPA also demonstrated the disturbance of biological processes associated with neuroinflammation and glutamatergic signaling (Figure 5f). Glutamatergic neurotransmission is the main excitatory stimulus in the central nervous system (CNS), and the binding of glutamate to its ionotropic glutamate receptors (iGluRs) is crucial for maintaining synaptic plasticity and neuronal survival. Recently, studies have demonstrated that AD patients exhibit excessive activity of extrasynaptic N-methyl-d-aspartate receptors (NMDARs), a type of iGluRs. This activity may result in neurodegeneration due to pronounced excitotoxicity caused by CREB inactivation and up-regulation of pro-death transcription factors like FOXO (64). This mechanism of neurodegeneration is further substantiated by the favorable clinical outcomes of memantine, an NMDAR antagonist currently administered for moderate to severe AD cases. Importantly, TAIPA findings also emphasize the impairment of synaptic structures (Figure 5f), a recognized hallmark of AD synaptopathology that associates with neurodegeneration. Specifically, this analysis indicates that the overexpression of miR-410-3p, miR-301a-3p, miR-320a-3p, miR-29a-3p, miR-181c-3p, miR-29c-3p, miR-660-5p is significantly associated with reduced levels of neurexin 3 (NRXN3), which encodes a crucial synaptic protein responsible for maintaining neural connections. The reduction in NRXN3 expression has been previously observed in individuals with AD (65), whereby the insufficiency of this factor could affect synaptic signaling and development. Overall, dysregulation of this miRNA-mRNA network may contribute to the impairment of synaptic plasticity, which is a critical component in cognitive processing. Lastly, MIRit analysis reveals the perturbation of the Wnt signaling pathway (Figure 5f), which has been previously reported in AD (66). In particular, our study found that an increase in miR-320a-3p levels may be responsible for the suppression of SERPINF1, also known as pigment epithelium derived factor (PEDF), a key neurotrophic protein implicated in AD development. In this concern, Huang et al. demonstrated that the downregulation of SERPINF1 in AD patients leads to diminished inhibition of the c-Jun N-terminal kinase (JNK) pathway, which in turn induces the expression of presenilin-1 (PS1), one of four proteins in the *γ*-secretase complex that cleaves amyloid precursor protein (APP) to produce *Aβ*_1*−*42_ fragments (67). In this context, increased levels of PS1 have been shown to induce the generation of *Aβ*_1*−*42_ fibrils (68), which eventually form neuritic plaques. In conclusion, MIRit’s analysis suggests that excessive transcription of miR-320a-3p may be responsible for the reduced expression of SERPINF1, resulting in the upregulation of PS1, which ultimately increases the synthesis of *Aβ*_1*−*42_ fragments. In the future, however, it will be imperative to undertake functional validation of these miRNAs, in order to comprehensively evaluate their role in AD pathogenesis and treatment.

## Conclusion

MiRNAs are essential biomolecules that regulate gene expression, and their dysregulation is frequently linked to various diseases. Since their discovery, there has been a significant effort to functionally characterize and describe miRNAs in pathological conditions. Although omic technologies, such as microarray and miRNA-Seq, provide us with the ability to measure expression variations of small transcripts, identifying their roles in disease mechanisms remains an unsolved problem. Specifically, current strategies for conducting integrative miRNA-mRNA analyses tend to produce limited success and are only applicable to datasets where the same individuals have been profiled for both miRNAs and mRNAs. For a detailed review of the tools available for integrative analysis of miRNAs and genes, please refer to Supplementary Table 9. Moreover, even after identifying significant miRNA-target pairs, it is not possible to thoroughly investigate the functional impact of perturbed miRNA-target interactions on biological pathways. To overcome these limitations, we have developed MIRit, an all-in-one R package that allows comprehensive analysis of miRNA and mRNA expression data, leading to the identification of compromised regulatory networks underlying pathological processes. Specifically, MIRit starts by identifying differentially expressed miRNAs and genes using the most established models in the field and then retrieves predicted and validated targets using accurate and up-to-date tools. Later, the integration of miRNA and gene expression is performed to identify influential interactions in which miRNA expression is negatively associated with that of its targets. To achieve this, paired datasets undergo correlation analysis, while unpaired datasets can be subjected to association tests and rotation gene set tests, making MIRit suitable also for meta-analysis. Instead of relying on all putative miRNA-target pairs, this data-driven approach selectively keeps miRNA-target interactions only when inverse relationships are detected in expression data, thereby reducing the number of false positive results. Lastly, MIRit introduces TAIPA, an innovative method for integrative miRNA-mRNA pathway analyses that considers the topological structure of biological pathways to identify the most disrupted molecular networks.

Here we described the performance of our proposed pipeline using two distinct datasets. The first study involved the investigation of altered miRNA-mRNA networks in the context of PTC samples, as compared to normal thyroid. In this case, MIRit analysis indicated a significant increase in miR-146b-5p and miR-146b-3p, resulting in reduced PAX8 expression and consequently decreased transcription of TG. This altered network may be responsible for the development of radioactive-iodine resistance in PTC patients, as reduced TG synthesis is responsible for decreased uptake and incorporation of radioactive iodine in cancer cells. Furthermore, we showcased the usage of MIRit with non-matched miRNA and mRNA expression data obtained from individuals with AD, compared to age-matched control samples. In this regard, MIRit has demonstrated the central role of miRNA dysregulation in various AD-related processes, such as neuroinflammation, glutamatergic signaling, and synaptic degeneration. Moreover, we postulate that the upregulation of miR-320a-3p may account for the downregulation of SERPINF1, which in turn causes overexpression of PS1, and thus excessive synthesis of *Aβ*_1*−*42_ fragments. In both of the provided case studies, the compromised networks identified by MIRit display significant relevance to the mechanisms of the disease, further confirming the suitability of MIRit for exploring perturbed miRNA-mRNA networks in multiple scenarios. In conclusion, MIRit proves to be a valuable tool for researchers investigating the effects of perturbed molecular networks in the context of integrative multi-omic miRNA-mRNA analyses, allowing users to thoroughly assess the disturbed mechanisms and functionally characterize the relative biological consequences. MIRit is completely open-source and freely available on GitHub.

## Availability and requirements

**Project name**: MIRit

**Project home page**: jacopo-ronchi/MIRit https://github.com/

**Operating system(s)**: Platform independent

**Programming language**: R, C++

**Other requirements**: Bioconductor

**License**: GPL v3.0

**Any restrictions to use by non-academics**: none

### List of abbreviations

AD: Alzheimer’s disease
APP: amyloid precursor protein
BA9: Brodmann area 9
BFS: breadth-first search
CAMERA: Correlation Adjusted MEan RAnk gene set test
CNS: central nervous system
DE-miRNAs: differentially expressed miRNAs
DEGs: differentially expressed genes
DIO2: iodothyronine deiodinase 2
DLB: dementia with Lewy bodies
FCS: functional class scoring
GEO: Gene Expression Omnibus
GO: Gene Ontology
GSEA: gene set enrichment analysis
HCC: hepatocellular carcinoma
iGluRs: ionotropic glutamate receptors
JNK: c-Jun N-terminal kinase
MDS: multidimensional scaling
mirDIP: microRNA Data Integration Portal
miRNA: microRNA
NMDAR: N-methyl-d-aspartate receptor
NRXN3: neurexin 3
ORA: over-representation analysis
PA: pathway analysis
PAX8: Paired Box 8
PDD: Parkinson’s disease dementia
PEDF: pigment epithelium derived factor
PS1: presenilin-1
PTC: papillary thyroid carcinoma
RMA: Robust Multi-array Average
SLE: systemic lupus erythematosus
SNP: single nucleotide polymorphism
SRA: Sequence Read Archive
TAIPA: Topology-Aware Integrative Pathway Analysis
TG: thyroglobulin
TPD: tangle-predominant dementia
TPO: thyroid peroxidase

## Declarations

### Ethics approval and consent to participate

This article did not involve any studies with human participants or animals performed by the authors.

### Consent for publication

Not applicable

## Availability of data and materials

MIRit source code is available on GitHub at: https://github.com/jacopo-ronchi/MIRit. The thyroid cancer and the Alzheimer’s disease datasets are available on GEO, accession numbers GSE63511, GSE63501, and GSE150696. Moreover, the files used to analyze both the datasets are available at: https://github.com/jacopo-ronchi/MIRit_supporting_files.

## Authors’ contributions

J.R. conducted the bioinformatic analyses, processed the data, developed the data analysis framework, and wrote the manuscript. M.F. provided funding for the project and critically reviewed the manuscript. Both authors have contributed substantially, directly, and intellectually to this work and have approved it for publication.

## Acknowledgements

We thank the authors of the thyroid cancer and Alzheimer’s disease datasets for sharing expression data in public databases.

## Supplementary material

### Supplementary Table 1

- File name: supplementary-table-1.csv
- File format: comma-separated values (CSV)
- Title: Supported species
- Description: a table summarizing the species supported by the MIRit pipeline

### Supplementary Table 2

- File name: supplementary-table-2.csv
- File format: comma-separated values (CSV)
- Title: Differentially expressed genes in PTC
- Description: a table listing differentially expressed genes in PTC samples compared to normal thyroid

### Supplementary Table 3

- File name: supplementary-table-3.csv
- File format: comma-separated values (CSV)
- Title: Differentially expressed miRNAs in PTC
- Description: a table listing differentially expressed miRNAs in PTC samples compared to normal thyroid

### Supplementary Table 4

- File name: supplementary-table-4.csv
- File format: comma-separated values (CSV)
- Title: miRNA-mRNA correlation in thyroid cancer
- Description: a table displaying the results of Spearman’s correlation analysis between miRNA and target expression

### Supplementary Table 5

- File name: supplementary-table-5.csv
- File format: comma-separated values (CSV)
- Title: Sequenced reads summary for GSE63501
- Description: a table reporting a summary of GSE63501 preprocessing

### Supplementary Table 6

- File name: supplementary-table-6.csv
- File format: comma-separated values (CSV)
- Title: Differentially expressed miRNAs in Alzheimer’s disease
- Description: a table listing differentially expressed miRNAs in AD patients compared to healthy donors

### Supplementary Table 7

- File name: supplementary-table-7.csv
- File format: comma-separated values (CSV)
- Title: Differentially expressed genes in Alzheimer’s disease
- Description: a table listing differentially expressed genes in AD patients compared to healthy donors

### Supplementary Table 8

- File name: supplementary-table-8.csv
- File format: comma-separated values (CSV)
- Title: inverse miRNA-mRNA associations in AD
- Description: a table listing the negative associations between differentially expressed genes and differentially expressed miRNAs in AD patients, as detected by Boschloo’s exact test

### Supplementary Table 9

- File name: supplementary-table-9.csv
- File format: comma-separated values (CSV)
- Title: description of currently available tools for integrative miRNA anlyses
- Description: a table listing the features of three available tools that are commonly used to perform integrative miRNA-mRNA analyses

