## Supplementary figures and images for "MIRit: an integrative R framework for the identification of impaired miRNA-mRNA regulatory networks in complex diseases"

### Supplementary Figure 1

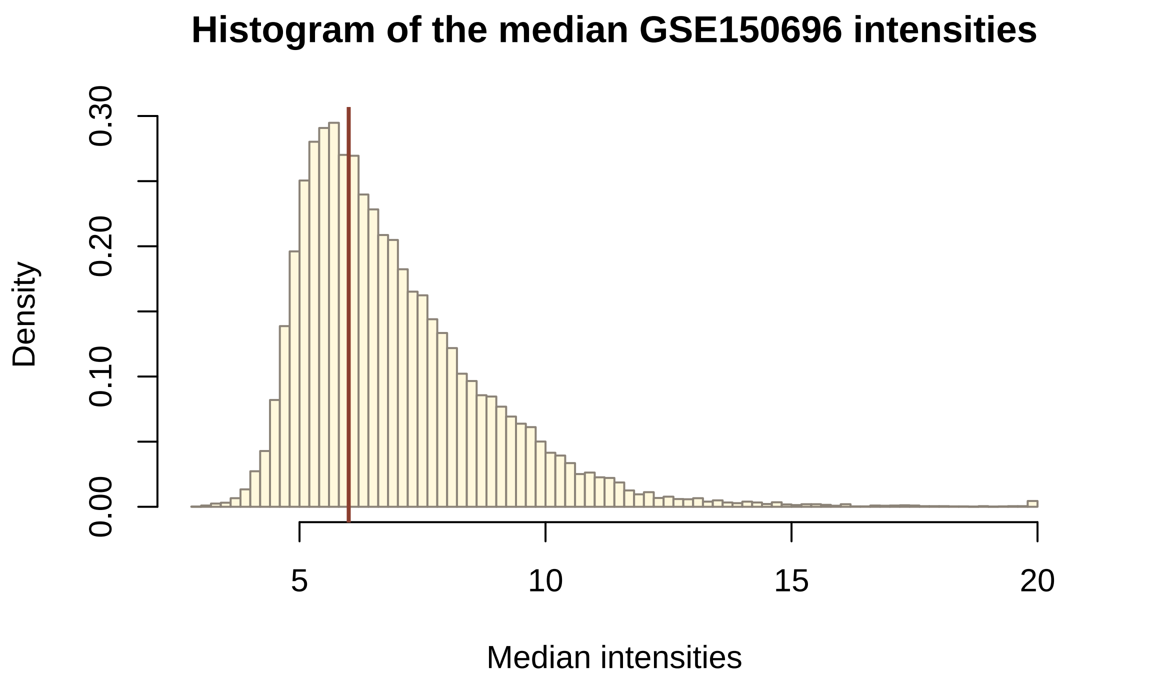
